# A human microbiota-associated mouse model of early-life malnutrition reveals persistent microbiome immaturity and limited response to fecal viral transplantation

**DOI:** 10.64898/2026.08.26.747040

**Authors:** Michael Shamash, Laura Carolina Camelo Valera, Corinne F. Maurice

## Abstract

Malnutrition is a leading cause of child mortality worldwide and has long-lasting health and socio-economic consequences. Studies have established causal links between the gut microbiota and childhood malnutrition, with key microbial signatures including delayed microbiome development and an enrichment of bacterial pathogens. While current dietary interventions improve growth and developmental outcomes, post-therapy regression to an immature microbial state is common. Fecal virome transplants (FVTs) represent a promising approach to reshape gut microbial communities, yet their therapeutic potential in early life remains poorly described. In this work, we established a diet-inducible human microbiota-associated (HMA) mouse model of early-life stunting, where malnourished pups were 35% lighter and 25% shorter than healthy controls. We developed a predictive model to quantify gut bacteriome development, identifying *Enterococcus* and *Clostridium* as primary drivers of healthy maturation. Our model revealed that the malnourished HMA mouse gut remains significantly immature compared to healthy controls and decoupled from the mouse’s chronological age. While a successful FVT from a healthy donor induced targeted changes in specific bacterial taxa, including a significant increase in *Enterococcus* species, it did not rescue physical growth or lead to broad community-level shifts. In contrast, a failed FVT from a different healthy donor revealed a significant mismatch between the donor virome and recipient bacteriome, indicating niche incompatibility that limits FVT efficacy.

Our work establishes a robust human microbiota-associated mouse model for studying maturation of the gut in early life, suggesting that FVT alone is insufficient to reproducibly reshape the malnourished gut. These findings highlight the need for synergistic strategies, combining viral interventions with nutritional supplementation for maximum therapeutic effect.

## Introduction

Child undernutrition remains a global health challenge, presenting primarily as stunting (height-for-age z-score < -2 standard deviations from the World Health Organization [WHO] median), and wasting (weight-for-height z-score < -2 standard deviations from the WHO median) (1). Together, these conditions affect over 30% of children worldwide and are responsible for 45% of all deaths in children under the age of five (2, 3). Recent research has shown that the gut microbiome of children with severe acute malnutrition follows a delayed development compared to healthy children (4–6). This microbial immaturity is characterized by low relative abundances in beneficial *Lactobacillaceae* & *Bifidobacteriaceae*, and an enrichment of opportunistic pathogens, such as *Enterobacteriaceae*. Considering that the defined succession of the gut microbiota in the first 1,000 days of life is critical for immune and physical development, these disturbances can lead to long-lasting health conditions, such as autoimmune and allergic diseases, as well as stunted growth (4, 6–12). While dietary interventions can improve the clinical symptoms of undernutrition in children, their gut microbiome halts development upon the cessation of therapy (4). Thus, there is an urgent need for therapies that promote the lasting maturation of the gut microbiome.

Bacteriophages (or phages) are bacteria-infecting viruses and major members of the human gut virome, the viral fraction of the microbiota. Phages are extensively described as regulators of bacterial community structure, metabolism, and evolution (13–17). While the healthy infant gut virome assembles rapidly and reproducibly, this process is disrupted in children with malnutrition, who exhibit altered viral diversity and composition compared to healthy controls (5, 18). Previous work from our lab demonstrated that *in vitro* cross-infections with phages and bacteria from healthy and stunted children, respectively, could shift bacterial community composition towards a healthy state (12). In addition, recent exciting studies show that fecal virome transplants (FVT) can remodel bacterial communities in contexts ranging from inflammatory bowel diseases to necrotizing enterocolitis (19–22), highlighting the therapeutic potential of this approach. During the first three years of life, the gut microbiome is highly dynamic and characterized by active kill-the-winner interactions with high abundances of lytic phages (18, 23, 24). Given this state of viral-driven community turnover, we hypothesized that early life represents an opportune window for FVT-mediated remodelling to restore healthy gut maturation trajectories in the context of undernutrition.

Existing mouse models of malnutrition initiate dietary insufficiencies only after weaning, leaving the dynamic pre-weaning developmental window largely unexplored (25–28). To address this gap and test our hypothesis, we first developed and validated an animal model of early-life malnutrition in human microbiota-associated mice, adapting a diet-inducible framework originally used in conventional adult mice (25). By colonizing germ-free breeders with healthy or stunted infant fecal samples, we could evaluate the long-term effects of vertical transmission and nutritional deficiency in their offspring (pups). Central to our analysis, we developed a predictive microbiota age model to quantify the gut microbial immaturity induced by malnutrition.

Our diet-inducible model of early-life malnutrition led to significant physical stunting in mouse pups. Those raised on the protein- and fat-deficient diet were approximately 35% lighter and 25% shorter than healthy controls at the time of weaning. Despite the expected selective engraftment of the original infant microbial communities into germ-free breeder mice, we observed high rates of vertical transmission from dams to pups. To attempt a rescue of the stunted phenotype and reshape stunted microbial communities, we administered four doses of healthy donor-derived FVT to stunted pups around the time of weaning. Despite the intervention, pups remained physically stunted and exhibited highly variable, targeted taxonomic shifts of their gut microbiota rather than broad community-level changes. This limited impact suggests that viral engraftment and modulation of the gut bacteriome are constrained by both host availability and resource limitation, reflecting phage-host incompatibility and abiotic filtering. Consequently, FVT alone is insufficient to broadly restructure the gut bacteriome. Since neither nutritional intervention alone (4) nor FVT alone is sufficient in remodeling the malnourished gut, future studies should evaluate their combination to overcome the ecological constrains identified here.

## Methods

### Human fecal sample collection

Human fecal samples were collected with the approval of protocol A04-M27-15B from the McGill University Institutional Review Board and protocol PR-16001 from the International Centre for Diarrhoeal Disease Research, Bangladesh (icddr,b). Samples were collected from healthy and stunted infants in Dhaka, Bangladesh, in collaboration with the lab of Dr. Dinesh Mondal at the icddr,b. Parents of the participants provided informed written consent for the utilization of their samples and met specific inclusion criteria. To assign infants to either the healthy or stunted growth category, we used the World Health Organization-recommended height-for-age Z-scores (HAZ < -2) to define stunting and HAZ ≥ -2 to define healthy growth. Exclusion criteria: children with diarrhea, respiratory infection, antibiotic use, or nutritional supplementation at enrollment. Sample collection was delayed and recorded for transient illness. Once collected, samples were stored at - 70°C until being used.

### Animal studies

All animal work was performed under a protocol approved by the Comparative Medicine and Animal Resource Centre Facility Animal Care Committee (CMARC FACC) at McGill University (protocol # MCGL-7999) in accordance with the Canadian Council on Animal Care guidelines. Germ-free C57BL/6 mice were housed in Techniplast IsoCages at the Research Institute of the McGill University Health Centre (RI-MUHC). For Experiment #1, 5-week-old germ-free mice were obtained from the University of Calgary; for Experiments #2 and #3, germ-free mice were bred in-house.

Mice were maintained on either a control diet (CON diet, D09051102i) or an isocaloric fat- and protein-deficient diet (MAL diet, D14071001Bi), both from Research Diets Inc. (New Brunswick, NJ, USA). Diets were irradiated by the manufacturer to ensure sterility, and food and autoclaved water were provided to all mice *ad libitum*.

At approximately 7 weeks of age, each germ-free mouse was colonized with a fecal sample from either healthy or stunted infant donors. Donor feces were resuspended in sterile reduced phosphate-buffered saline (rPBS; 0.02 µm-filtered) at a ratio of 1 g feces to 15 mL rPBS. Each mouse received 200 µL of the resulting fecal slurry by oral gavage. Following a one-week stabilization period, breeding trios (two females, one male) were established. Donor assignment was fixed per cage, so that all mice within a breeding trio shared the same inoculum. Upon confirmation of pregnancy, breeding trios were split, and each female was moved to their own cage. Fecal sampling and body measurements of the dams (tail length, body length, body weight using digital calipers) were performed regularly. To minimize stress, sampling and measurements were paused from 1 week before pup birth until 1 week after pup birth.

Experiment #1 was an observational study in which mice with different donors were maintained on either CON or MAL diets, bred, and the pups monitored until six weeks of age (**Figure 1A**). Experiments #2 and #3 were intervention studies involving administration of fecal virome transplants (FVTs) to pups (**Figure 3A**).

**Figure 1.**
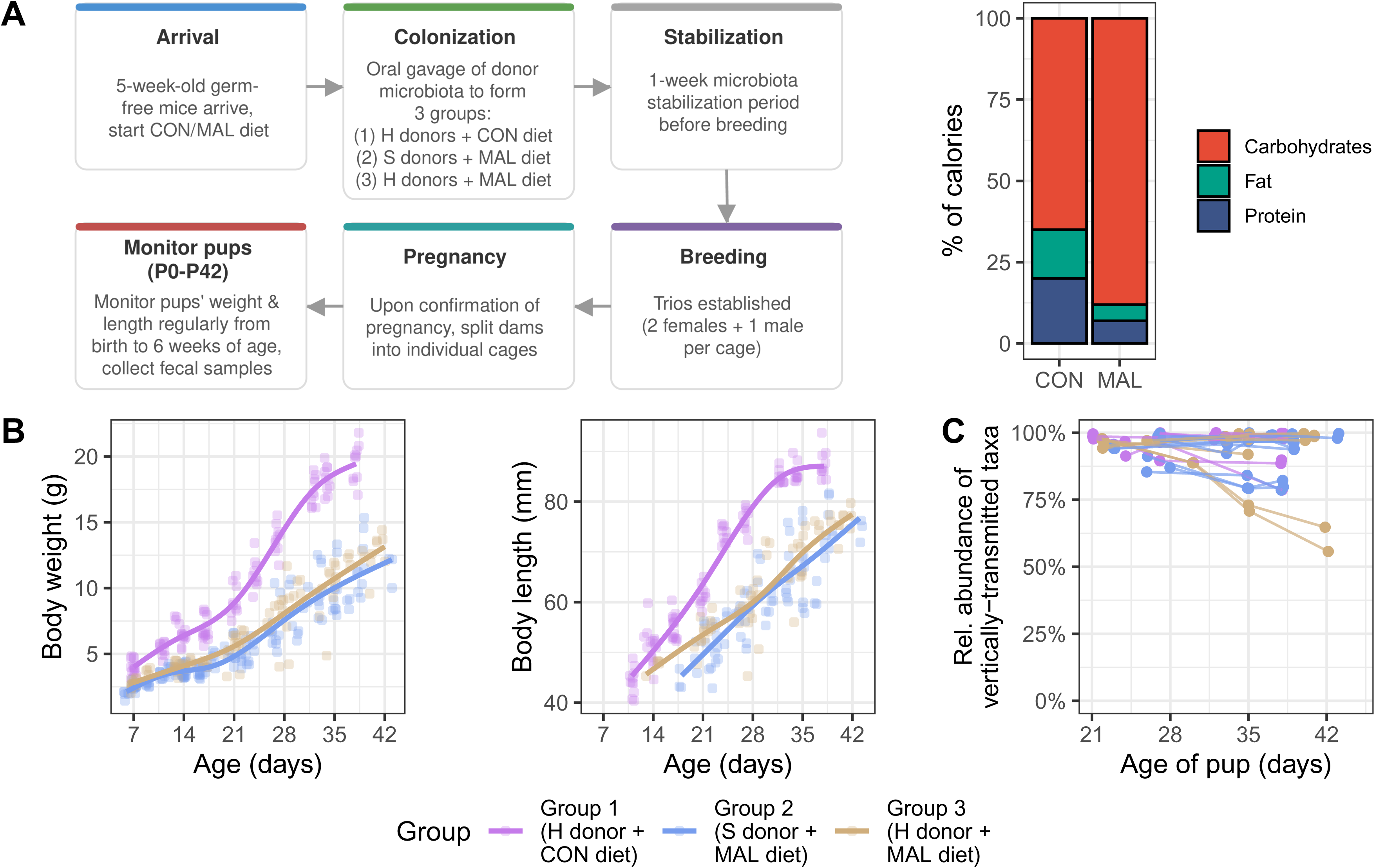
A protein- and fat-deficient diet induces physical stunting in human-microbiota-associated mouse pups. **(A)** Overview of the experimental design and dietary composition for the intergenerational observational study (Experiment #1). Five-week-old germ-free mice were placed on either a control (CON) or a malnourished (MAL) diet. The macronutrient compositions of both diets are shown (right). Mice were colonized via oral gavage with donor microbiota to establish three experimental groups: healthy (H) donors on a CON diet (Group 1), stunted (S) donors on a MAL diet (Group 2), and H donors on a MAL diet (Group 3). Following a one-week stabilization period for the microbiota, breeding trios (two females and one male per cage) were established. Once pregnancy was confirmed, the dams were split into individual cages. The pups were monitored regularly from birth to 42 days (6 weeks) of age to track weight and length, alongside the collection of fecal samples. **(B)** Absolute body weight and length of pups over time. **(C)** Dam-to-pup vertical transmission, represented by the cumulative relative abundance of a dam’s bacterial taxa detected in her resulting pups.

Live FVTs were prepared by resuspending healthy infant donor feces in rPBS (1 g/15 mL). The slurry was centrifuged at 1,000 *g* for 1 minute to remove large debris, followed by centrifugation of the supernatant at 10,000 *g* for 10 minutes to pellet bacterial cells. The final supernatant was passed through a 0.20 µm filter to recover the live FVT filtrate. Temperature-inactivated (TI) FVTs were prepared identically, followed by three rounds of thermal shock to denature phage particles (flash-freezing in liquid nitrogen for 3 minutes, then heating at 95 °C for 3 minutes). The TI-FVT was then treated with a DNase cocktail (100 U TURBO DNase, Thermo Fisher Scientific, Waltham, MA, USA; 1,000 U Benzonase, Sigma-Aldrich, Burlington, MA, USA) at 37 °C for 1 hour to remove residual DNA. The DNase was inactivated at 75 °C for 30 minutes. All FVTs were stored in Protein LoBind tubes on ice and administered within 2 hours of preparation. Pups received 40 µL of FVT (live, temperature-inactivated, or rPBS vehicle) via oral gavage upon reaching a body weight of 10 g (approximately postnatal day 28). Pups received two doses prior to weaning and two additional doses post-weaning, for a total of four doses over a 10-day period (averaging 2 days between doses). Pups were monitored for one month following the final dose.

### Bacteriome sequencing and bioinformatic analysis

Genomic DNA was extracted from human and mouse fecal samples using the QIAamp PowerFecal Pro DNA kit (QIAGEN, Hilden, Germany) according to the manufacturer’s instructions.

In Experiment #1, the 16S rRNA gene was amplified using universal 27F (5’-AGRGTTYGATYMTGGCTCAG-3’) and 1492R (5’-CGGYTACCTTGTTACGACTT-3’) primers with Invitrogen Platinum SuperFi PCR Master Mix (Thermo Fisher Scientific, Waltham, MA, USA). Amplicons were barcoded, pooled, and prepared for sequencing using the Native Barcoding 96 V14 kit (SQK-NBD114.96; Oxford Nanopore Technologies [ONT], Oxford, UK). In Experiments #2 and #3, a modified high-throughput library preparation approach was implemented (29). The 16S rRNA gene was amplified using M13-tailed, barcoded 27F and 1492R primers (5’-GTAAAACGACGGCCAGTG-[8bp barcode]-[27F/1492R primer sequence]-3’) and Invitrogen Platinum SuperFi PCR Master Mix. Following equimolar pooling of amplicons and purification, a DBCO-tagged M13 primer with phosphoramidite spacers (5’-/5DBCOTEG/GCTTGGGTGTTTAACC/iSpC3//iSpC3//iSpC3//iSpC3/GTAAAACGACGGCC AGTG-3’) was used in a second PCR round to label amplicons at the 5’ and 3’ ends. The final DBCO-labelled pool was purified and ligated to motor proteins using the Rapid Adapter Auxiliary V14 kit (EXP-RAA114, ONT). All libraries were sequenced on a MinION Mk1B instrument using R10.4.1 flow cells (ONT). Basecalling and demultiplexing were performed using Dorado (v0.8.3) (https://github.com/nanoporetech/dorado) with the dna_r10.4.1_e8.2_400bps_sup@v5.0.0 model. Reads were filtered by length to retain sequences between 1.3 kb and 1.7 kb (target 1.5 kb ± 0.2 kb). Taxonomic identification was performed using Emu (v3.5.2) (30), and downstream diversity and composition analyses were conducted in R (v4.5.1) using phyloseq (v1.52.0) (31) and vegan (v2.7-2) (32).

### Bacteriome age prediction and maturation modelling

To model the relationship between bacteriome composition and chronological age in Experiment #1, we used a supervised machine learning approach. First, dimensionality reduction was performed via Principal Coordinates Analysis (PCoA) based on weighted UniFrac distances. The resulting PCoA eigenvectors (axes) were extracted to serve as predictors for partial least squares regression (PLSR), using the pls package (v2.8-5) (33). The PLSR model was trained exclusively on pup samples from Group 1 (healthy donor background, control diet) to define a baseline maturation trajectory. The optimal number of latent components was determined by minimizing the root mean square error of prediction (RMSEP). To assess the impact nutritional deficiency, this trained model was then used to predict the “microbiota age” of pups from Group 3 (same healthy donor background, deficient diet).

To evaluate differences in predicted age trajectories between groups, we used generalized additive mixed models (GAMMs) via the mgcv package (v1.9-3) (34). The models were fit using the formula: age_predicted ∼ s(age_actual, by = group) + group. Statistical differences between group maturation curves were assessed using the emmeans package (v2.0.0) (35).

Bacterial taxa driving the PLSR model were identified by integrating regression coefficients with taxonomic loadings. Specifically, the PLSR coefficients for each PCoA axis were multiplied by the original taxonomic biplot coordinates. The importance of each genus was ranked by its Euclidian distance from the origin in ordination space, highlighting the taxa most strongly associated with the maturation process. A heatmap of the most important genera was generated using ComplexHeatmap (v2.24.1) (36) to visualize mean abundance patterns over the first six weeks of life. For each genus, a Pearson correlation coefficient (*r*) was calculated against pup age (in days). These values were squared to obtain the coefficient of determination (R^2^), representing the proportion of age variance explained by that genus. We calculated a signed R^2^ by multiplying the R^2^ value by the sign of *r* to indicate both the strength and direction of the association with chronological age.

### Virome sequencing and bioinformatic analysis

Fecal viral DNA extraction and library preparation were performed as previously described (37). Equimolar pools were sequenced on an Illumina NovaSeq X Plus instrument using 150 bp paired-end reads (SeqCenter, Pittsburgh, PA, USA).

Raw reads were trimmed and filtered using fastp (v0.23.4) (38) with the following parameters: --detect_adapter_for_pe -q 15 --cut_right --cut_window_size 4 --cut_mean_quality 20 --length_required 100. To remove host contamination, trimmed reads were mapped against the *Homo sapiens* (GRCh38) and *Mus musculus* (GRCm39) genomes using bowtie2 (v2.5.2) (39). Decontaminated reads from each sample were individually assembled *de novo* using metaSPAdes (v3.15.4) (40, 41) in --meta mode. Viral contigs were identified from the resulting assemblies using geNomad (v1.7.4) (42). Only viral contigs ≥ 1 kb were retained for downstream analysis. Viral contigs were dereplicated into viral operational taxonomic units (vOTUs) using BLASTN (v2.14.0) (43) and the anicalc.py and aniclust.py scripts from CheckV (44). Clustering was performed using a threshold of 95% average nucleotide identity (ANI) over 85% alignment fraction (AF) of the shorter contig. To quantify vOTU abundance, decontaminated reads were mapped back to the dereplicated vOTUs using bowtie2 (v2.5.2). Coverage statistics were generated using the ‘samtools coverage’ command (v1.18) (45). A vOTU was considered present in a sample only if it met the detection threshold: mean depth of coverage ≥1X and breadth of coverage ≥75% (46). Viral hosts were computationally predicted using iPHoP (v1.0.0, Sept_2021_pub database) (47), and viral lifestyles (temperate or non-temperate) were predicted using BACPHLIP (v0.9.6) (48). Diversity and community composition analyses were performed in R (v4.5.1) using the phyloseq (v1.52.0) and vegan (v2.7-2) packages.

## Statistical analyses

To model pup body measures over time, GAMMs were implemented in R (v4.5.1) (49) using the mgcv package (v1.9-3) (34). The following formula was used: weight/length ∼ s(age, by = group) + group + s(litter, bs = “re”) + s(litter, age, bs = “re”). The emmeans package (v2.0.0) (35) was used to evaluate pairwise differences between groups at specific timepoints. Visualization of the fitted smooth terms was performed using the gratia package (v0.11.1) (50).

Bacteriome and virome alpha diversity metrics (observed richness and Shannon index) were modelled with a GAMM with the following equation: y ∼ group + s(age_relative, by = group) + s(animalID, bs = “re”) + s(damID, bs = “re”), where age_relative corresponds to the pup age in days relative to the first FVT dose. The s(damID, bs = “re”) term was not included for Experiment #3 models as there was only one breeding dam per experimental group.

Differentially abundant bacterial taxa and vOTUs were identified using MaAsLin 3 (v1.0.2) (51) with a significance threshold of q < 0.1, log transformation, and no normalization (inputs were already relative abundances, also known as total sum scaled). The following formula was used: ∼ Timepoint + (1 | AnimalID), where Timepoint is a 2-level factor corresponding to either before or after FVT. To isolate the specific effects of the FVT, bacterial taxa/vOTUs were excluded from the final results if they also showed significant differential abundance (and in the same direction) in the TI-FVT or rPBS vehicle control groups.

## Results

### A protein- and fat-deficient diet induces physical stunting in mouse pups

To evaluate the impact of maternal and early-life nutrition on gut microbiome assembly and growth, germ-free adult mice were colonized with fecal samples from two healthy (Donors 1 [16-month-old male] and 2 [17-month-old female]) and two stunted (Donors 3 [16-month-old male] and 4 [17-month-old female]) infant donors (Experiment #1, **Figure 1A**). Breeders and their pups were maintained on either a control (CON) or an isocaloric fat- and protein-deficient (MAL) diet (**Figure 1A**). Across 9 litters, we monitored a total of 30 pups (Group 1, n = 10; Group 2, n = 12; Group 3, n = 8). Body weight and measurements (tail and body length) were collected beginning around postnatal day (P)7 and P14, respectively. Fecal samples were collected from pups as soon as they began producing feces, around P21.

Pup weight, body length, and tail length were significantly associated with age (*p* < 0.001, generalized additive mixed model), with the MAL diet inducing a consistent stunting phenotype. By weaning (P28), pups on the MAL diet were approximately 35% lighter and had 25% shorter body and tail lengths compared to those on the CON diet (**Figure 1B**, **Supplementary Figure 1A**). These effects were significant across all timepoints, regardless of donor inoculum (estimated marginal means contrasts *p* < 0.05).

We next evaluated the colonization efficiency of the human stool samples into the germ-free dams and the subsequent vertical transmission to their pups. Consistent with previous human microbiota-associated mouse models (52–54), we observed a bottleneck during human-to-mouse engraftment. In the dams, bacterial taxa (species level) derived from the human inoculum accounted for a mean of 27.5% of total bacterial relative abundance. Despite this initial bottleneck, vertical transmission from dam to pup was highly efficient across all groups. An average of 94.8% of the pup bacteriome was composed of taxa originating from the dam (**Figure 1C**). Virome sequencing data were unavailable for this experiment due to technical challenges during library preparation, specifically poor amplification yielding DNA concentrations too low for sequencing.

### The malnourished gut bacteriome is microbially immature

We next evaluated the impact of the MAL diet on the development of the gut bacteriome in pups. We first modelled alpha diversity metrics (observed richness and Shannon index) as a function of diet and donor health state (**Figure 2A**). While bacterial richness was not significantly associated with pup age in any group (global smooth *p* = 0.99), the Shannon index increased significantly over time across all groups (global smooth *p* < 0.001). Group 2 (stunted donors, MAL diet) richness was significantly higher than Group 1 (healthy donor, CON diet) between P20 and P36, and higher than Group 3 (healthy donors, MAL diet) between P20 and P40. Similarly, the Shannon index for Group 2 was significantly higher than Group 1 from P20 to P38, but lower than Group 3 from P20 to P28. Group 3 Shannon index also surpassed Group 1 after P38 (**Figure 2A**).

**Figure 2.**
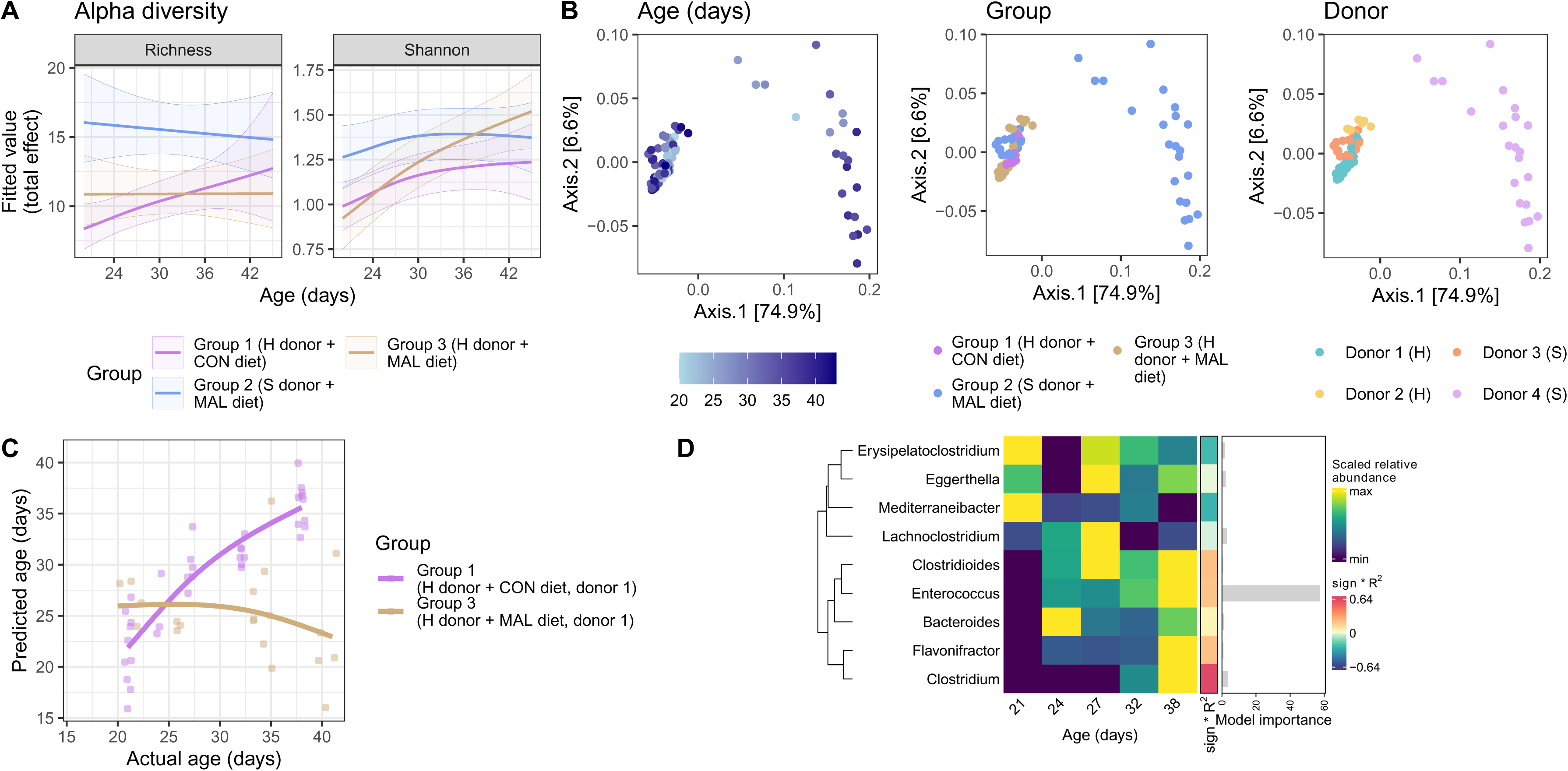
Stunted pups have immature gut bacterial communities. **(A)** Modelled alpha diversity trajectories using generalized additive mixed models (GAMMs). The fitted smooths represent the relationship between each metric and pup age (in days), with shaded ribbons indicating 95% confidence intervals. **(B)** Bacterial community structure visualized via principal coordinates analysis (PCoA) of weighted UniFrac distances. Points are colored by pup age (left), experimental group (centre), and donor ID (right). **(C)** Partial least squares regression model performance for predicting pup chronological age from bacteriome composition. Predicted age versus actual age (in days) is shown for both training (Group 1; healthy donor, CON diet) and testing (Group 3; healthy DONOR, MAL diet) datasets. **(D)** Heatmap of the most important bacterial genera in the age predictor model, with mean relative abundance in Group 1 pups over time (min/max scaled for each genus). Accompanying coloured strip and bar plot indicate relative overall contribution and importance of each genus to the model, respectively.

To evaluate community-level relatedness, we computed beta diversity using weighted UniFrac distances (**Figure 2B**). While sequencing depth (*p* = 0.04, R^2^ = 0.0184) and pup age (*p* = 0.001, R^2^ = 0.0212) were significant drivers of composition, donor ID was the primary determinant of community structure (*p* = 0.001) and explained 74.3% of the total variance (PERMANOVA test). Given that severe acute malnutrition is associated with delayed gut bacteriome maturation in infants, and that age was a significant factor in our community analysis, we sought to determine if the MAL diet induced a microbially immature phenotype in pups. To account for the strong effect of donor ID, we trained a partial least squares regression (PLSR) model exclusively on Group 1 pups (healthy donor background, CON diet) to establish a baseline trajectory for gut bacteriome maturation. The model demonstrated high accuracy on training data, with a root mean square error (RMSE) of 2.97 days and R^2^ = 0.803 (**Figure 2C**). When this baseline model was used to predict the microbiota age of Group 3 pups (same healthy donor, but MAL diet), we observed a distinct shift in development. While early timepoints (P20 to P22) revealed predicted microbiota ages that exceeded the pups’ chronological age, this early maturation was not sustained. Over time, the predicted microbiota age decreased significantly, lagging the chronological age of the animals after P26, supporting microbial immaturity (estimated marginal means contrasts *p* < 0.01). We limited this analysis to the healthy donor background, as donor ID was the dominant driver of community structure (74.3% of variance; **Figure 2B**). Since the PLSR predictors are derived from this ordination space, applying the Group 1-trained model to stunted-donor pups (Group 2) would conflate donor origin with maturation rather than isolate the effect of diet.

Finally, we identified the specific bacterial genera driving the age-prediction model in Group 1 pups (**Figure 2D**). Taxa were ranked by their importance to the model and the direction of their association with age was determined. *Enterococcus* and *Clostridium* were identified as the most age-discriminatory taxa, both of which increased in relative abundance with age. In contrast, *Lachnoclostridium* exhibited a transient increase followed by a decline as the pups aged.

### Fecal virome transplants have limited impact on community-level microbiome structure in early-life malnutrition

Following the observation that the MAL diet delays bacteriome development, we next explored if fecal virome transplants (FVTs) from healthy donors could reshape the gut microbiota of stunted pups. The experimental designs for Experiments #2 and #3 had the same longitudinal structure of Experiment #1, with the addition of four healthy donor-derived FVT doses administered via oral gavage around the time of weaning (**Figure 3A**). In Experiment #2, the same healthy and stunted donor material from Experiment #1 was used (Donors 1 and 3, respectively). For Experiment 3, a new pair of 14-month-old healthy and stunted male donors was used. Control groups received either temperature-inactivated FVT (TI-FVT) or a vehicle control (rPBS). However, we report an unexpected loss of the litters allocated to the rPBS vehicle control in Experiment #2 prior to the FVT, which meant that this control could only be included in Experiment #3. No adverse clinical signs were observed following FVT administration.

**Figure 3.**
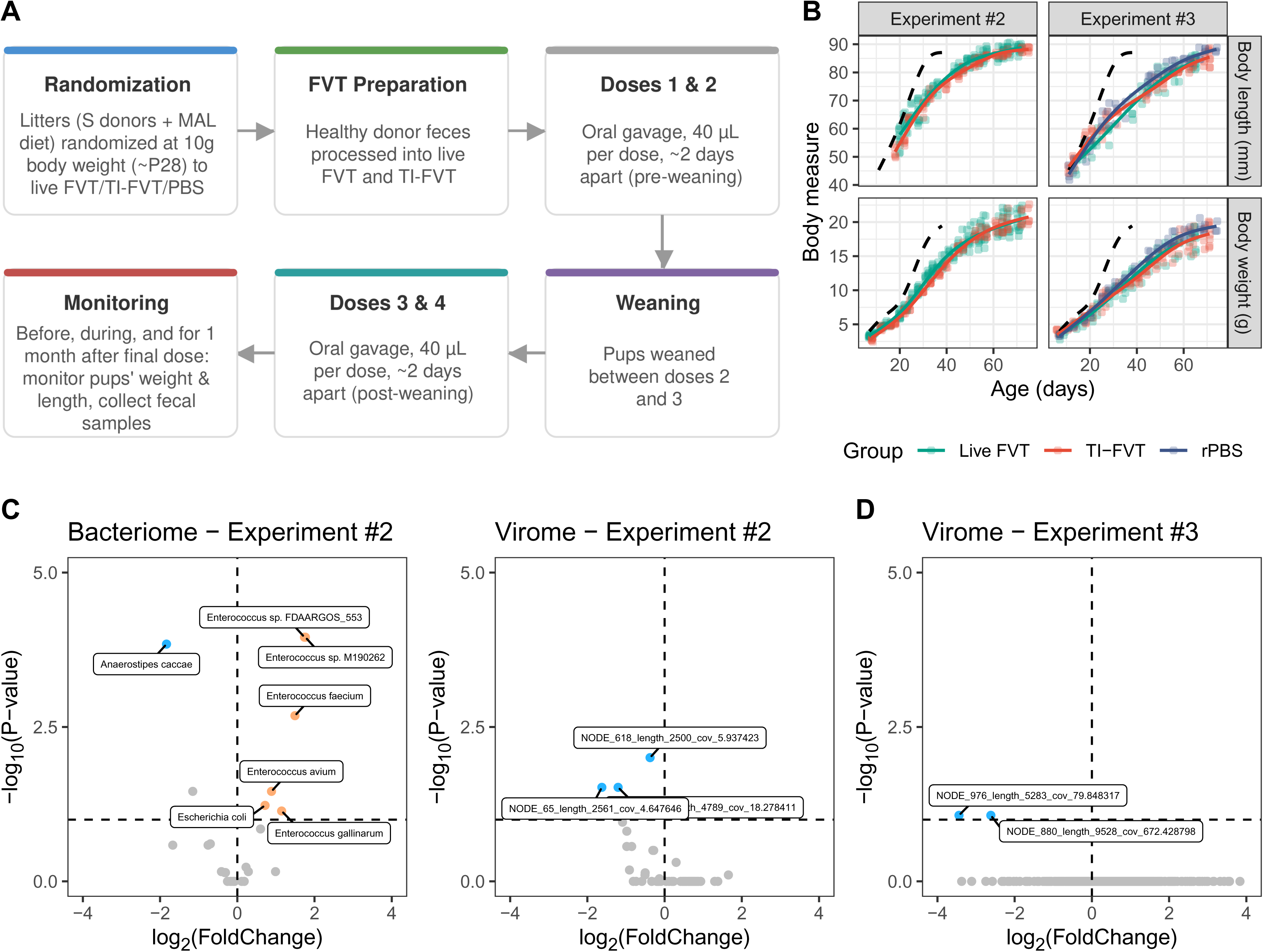
Fecal virome transplants exert limited effects on the gut microbiota of stunted pups. (**A)** Overview of the experimental design for the fecal virome transplant (FVT) intervention studies (Experiments #2 and #3). Litters with a stunted donor microbiota and on a malnourished diet (S donors + MAL diet) were randomized at approximately postnatal day 28 (P28; ∼10 g body weight) to receive live FVT, temperature-inactivated FVT (TI-FVT), or an rPBS vehicle control. Pups received a total of four 40 µL doses via oral gavage, spaced approximately two days apart: two pre-weaning doses and two post-weaning doses, with weaning occurring between the second and third dose. Pup weight, length, and fecal samples were monitored before, during, and for one month following the final dose. **(B)** Absolute body weight and length of pups over time. The dashed line represents the average trajectory of Group 1 pups (healthy donor background, CON diet) established in Experiment #1. **(C)** Volcano plots of differentially abundant bacterial taxa and vOTUs from Experiment #2 and **(D)** vOTUs from Experiment #3. Horizontal dotted line indicates the significance threshold of q = 0.1.

The FVT had no effect on the stunting phenotype in pups. Pups receiving live FVT, TI-FVT, or rPBS had near-identical growth dynamics in terms of in terms of body weight, body length, and tail length (**Figure 3B**, **Supplementary Figure 1B**). The growth of all groups remained stunted relative to the healthy control reference trajectory established in Experiment #1.

We next evaluated the broad response of the gut bacteriome and virome to FVT. In Experiment #2, alpha diversity metrics (observed richness and Shannon index) were unaffected by FVT (**Supplementary Figure 2A**). While viral communities in pups receiving live FVT exhibited a significant shift from their own baseline (PERMANOVA *p* = 0.029), their bacterial communities remained stable (*p* = 0.302). Importantly, by the end of the experiment, neither the viral (*p* = 0.64) nor the bacterial (*p* = 0.17) communities in the live FVT group differed significantly from those in the TI-FVT control group (**Supplementary Figure 2B, 2C**). Similarly, in Experiment #3, alpha diversity was unchanged by FVT relative to both control groups (**Supplementary Figure 2D**). At the whole community level, FVT had no significant effect; communities did not shift significantly from baseline, nor were there significant differences between the live FVT, TI-FVT, and rPBS groups by the end of the study (**Supplementary Figure 2E, 2F**).

Although the FVT did not induce broad community-level shifts, we investigated whether it exerted more targeted effects on specific taxa. We defined a bacterial species or viral operational taxonomic unit (vOTU) as responsive only if it was differentially abundant in the live FVT group and remained unchanged (or moved in the opposite direction) in the TI-FVT and rPBS control groups within the same experiment. In Experiment #2, several bacterial taxa had significant abundance shifts in response to FVT (**Figure 3C**). Specifically, *Escherichia coli* and five *Enterococcus* species increased in relative abundance post-FVT, while *Anaerostipes caccae* significantly decreased (**Supplementary Figure 3**). The virome response in Experiment #2 was characterized by three vOTUs that significantly decreased in abundance after FVT (**Figure 3C**, **Supplementary Figure 3**). These vOTUs were computationally predicted to infect *Bacteroides fragilis* and *Erysipelatoclostridium ramosum*. In Experiment #3, no bacterial taxa met our criteria for FVT responsiveness; however, two vOTUs significantly decreased in abundance following FVT. Both vOTUs were predicted to infect *Enterococcus* species (**Figure 3D**, **Supplementary Figure 3**).

### Limited host availability constrains FVT-mediated remodelling of bacterial communities

To investigate the limited impact of FVT on the pup bacteriome, we analyzed the overlap between the predicted bacterial hosts of the donor virome and the taxa present in the recipient pups at the time of FVT administration. We computationally predicted genus-level hosts for the vOTUs detected in the FVT (**Figure 4A**). In Experiment #2, 66.7% of FVT vOTUs (58 of 87) were assigned a host, representing 95% of the FVT’s total viral community abundance. Host assignment was lower in Experiment #3, where only 56.6% of FVT vOTUs (47 of 83) were assigned a host, accounting for just 11% of the community abundance. The most abundant vOTU in the Experiment #3 FVT (84% relative abundance) could not be assigned a host.

**Figure 4.**
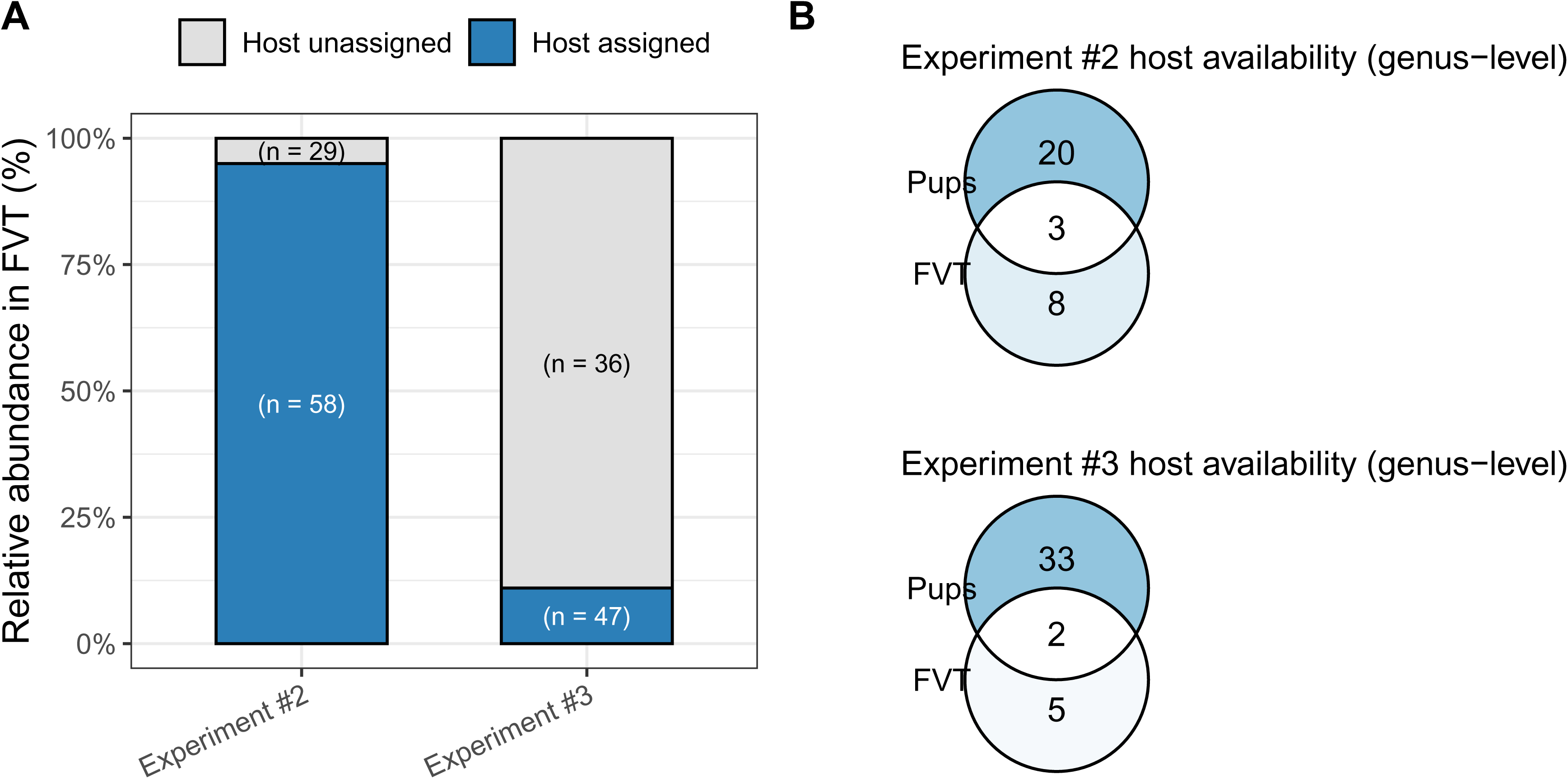
Limited host availability for fecal virome transplant bacteriophages. **(A)** Stacked bar charts displaying the relative community abundance of FVT vOTUs with computationally assigned versus unassigned bacterial hosts in Experiments #2 and #3. vOTU counts (n) are indicated within the respective bars. **(B)** Genus-level overlap between the predicted bacterial targets of the FVT and the bacteriome of recipient pups at baseline. In Experiment #2, shared genera include *Bacteroides*, *Bifidobacterium*, and *Clostridium*. In Experiment #3, shared genera include *Bacteroides* and *Bifidobacterium*.

In general, host availability was low across both FVT experiments (**Figure 4B**). In Experiment #2, where FVT led to changes in gut bacteriome composition, only 27.3% of the FVT’s predicted host genera (3 of 11) were present in pups at baseline, including *Bacteroides*, *Bifidobacterium*, and *Clostridium*. Similarly, in Experiment #3, where the bacteriome did not respond to FVT, only 28.6% of predicted host genera (2 of 7) were detected (*Bacteroides* and *Bifidobacterium*). Interestingly, the bacterial taxa that were significantly differentially abundant in Experiment #2, *Enterococcus* and *Anaerostipes*, had no vOTUs predicted as targeting them within the corresponding FVT.

Analysis of the predicted vOTU lifestyles within the FVT indicated a majority of the vOTUs were non-temperate. Across both experiments, approximately 90% of the vOTUs within the FVTs were non-temperate (Experiment #2: 90.8%, n = 79; Experiment #3: 88%, n = 73), with only a small fraction of the detected vOTUs carrying genes related to lysogeny (**Supplementary Figure 4**).

## Discussion

The gut microbiota follows a stepwise assembly over the first few years of life (24, 55). This assembly is delayed in the context of acute malnutrition (4). The aim of our study was to use fecal virome transplants to reshape the gut microbiota in the context of early-life undernutrition. To achieve this, we developed an animal model of early-life malnutrition, where germ-free breeder mice are first colonized with stunted infant donor fecal samples and maintained on either a control or isocaloric fat- and protein-deficient diet. Importantly, by using an intergenerational model, we were able to capture the pre-weaning period, often left unexplored by current models that only initiate dietary interventions after weaning (25–28). The strength of this model is further highlighted by the high heritability of the bacterial community observed from dams to pups.

After breeding the adult mice, we studied the physical growth and gut microbial composition of their pups until sexual maturity. Pups raised on a deficient diet were physically stunted and remained in a persistent state of microbial immaturity relative to healthy controls. A key finding of our age prediction model was the identification of *Enterococcus* as the most significant predictor of healthy development; its abundance being positively correlated with age in healthy pups. A recent meta-analysis of 3,154 gut metagenomes from 1,827 healthy infants revealed that *Enterococcus* abundance is negatively associated with age in healthy infants (24). This contrast suggests that our findings may be specific to the Dhaka cohort under study here, or alternatively, a result of using a human microbiota-associated animal model.

Given this state of gut microbial immaturity in stunted pups, we hypothesized that introducing healthy donor-derived phages via FVT could catalyze gut microbial community maturation. Although the FVT had limited effects at the community-level, it did lead to targeted effects on bacterial taxa in one of the two intervention experiments. In Experiment #2, we observed a targeted increase in five *Enterococcus* species and *Escherichia coli* following FVT. Interestingly, the two litters in this experiment responded with different magnitudes. The first litter to receive FVT had a greater increase in *Enterococcus* than the litter which was born two weeks later.

The limited community-level impact of FVT likely stems from multiple ecological constraints. Our host prediction analysis revealed significant mismatch between the donor virome and recipient bacteriome. In both experiments, fewer than 30% of predicted host genera were present in the pups at the time of transplant. Even when a host genus or species is present, strain-level specificity likely narrows the host range even further (56). While our full-length 16S rRNA marker gene sequencing successfully captures species-level dynamics, it cannot resolve strain-level variations that may further dictate phage susceptibility (57). It is also possible that the FVT led to strain switching within a species, which would not alter total taxonomic abundance but may have functional impacts (58, 59). Future studies with additional deep metagenomics sequencing will be essential to validate these strain-level dynamics.

Although we hypothesized that the active kill-the-winner dynamics typical of healthy early-life development would provide an opportune window for FVT-mediated remodelling (18, 23, 24), the limited efficacy of our intervention suggests that these viral interactions may be altered in the context of malnutrition. While our computational predictions confirmed that the administered healthy FVTs were mostly composed of non-temperate phages, they failed to induce widespread community changes. This resistance may be driven by the presence of bacterial lysogens in the recipient gut which could confer superinfection immunity against newly introduced phages (23, 60). To overcome this, future FVT applications should evaluate alternative windows of administration. Intervening earlier in the pups’ pre-weaning period or administering the FVT directly to the nursing dams may be necessary to reshape the successional trajectory of the pups’ gut microbiota.

Beyond top-down viral dynamics, the limited efficacy of the FVT highlights resource availability as a fundamental bottom-up constraint. Even if the introduced phages successfully targeted their bacterial hosts, opening new ecological niches, healthy microbial populations may not expand without additional essential macro- and micronutrients as pups were maintained strictly on the deficient MAL diet throughout the intervention (61, 62).

Finally, technical factors such as sample freshness may influence the FVT due to a loss of viral titer or infectivity after prolonged storage (21). While using fresh donor material is ideal, logistical constraints may make this impractical. Therefore, future FVT protocols should incorporate pre-intervention screening, such as quantifying virus-like particles (VLPs) and confirming infectivity via *in vitro* assays, to ensure the therapeutic viability of the transplanted virome.

Our work establishes a robust human microbiota-associated mouse model for studying the maturation of the malnourished gut in early life. Our findings suggest that FVT alone may be insufficient to reshape bacterial communities during early-life development. Future research should evaluate the synergistic potential of combining FVT with short-term nutritional interventions, such as peanut-based ready-to-use therapeutic food (RUTF) (11).

## Acknowledgements

This work was funded by a Canadian Institutes of Health Research Canada Project grant (PJT-175065), University of Toronto Canadian Wide Bacteriophage Therapy Acceleration Fund, and Canada Research Chair in Gut Microbial Interactions (Tier 2) to Corinne F. Maurice. Michael Shamash is supported by the Canadian Institutes of Health Research Canada Graduate Scholarship to Honor Nelson Mandela (CIHR CGS-D; #DF2-187718), and the *Fonds de recherche du Québec-Santé: Bourse de formation au doctorat* (FRQS; #311071). The authors would like to acknowledge Jackie Alford and Ezrah Isaac Roy for their help with preparing samples for sequencing, as well as Cynthia Faubert, Catherine Hudon, Frédérique Nadeau, and Stephanie Tang for their help with experiment planning and animal handling.

## Data and code availability

Code used for data analysis is available at https://github.com/mshamash/mouse_fvt_manuscript.

Sequencing data has been deposited to the NCBI SRA under accession number PRJNA1464576.

## Author contributions

M.S. conceived and performed the experiments, collected and processed samples for analysis, conceived and performed the analyses, prepared figures and tables, authored and reviewed drafts of the manuscript, and approved the final draft. L.C.C.V. performed the experiments, collected and processed samples for analysis, reviewed drafts of the manuscript, and approved the final draft. C.F.M. conceived the experiments and analyses, reviewed drafts of the manuscript, and approved the final draft.

## Declaration of generative AI and AI-assisted technologies in the writing process

During the preparation of this work the authors used Google Gemini (Alphabet Inc., Mountain View, USA) to improve sentence clarity and readability. Overall sentence meaning was not altered, and there were no changes to any of the presented data, facts, or conclusions by using this tool. After using this tool, the authors reviewed and edited the content as needed and take full responsibility for the content of the published article.

## Supplementary Figure Legends

**Supplementary Figure 1. Body length measurements of pups. (A)** Pup tail length measurements in Experiment #1 and **(B)** Experiments #2 and #3. The dashed line represents the average trajectory of Group 1 pups (healthy donor background, CON diet) established in Experiment #1.

**Supplementary Figure 2. Alpha and beta diversity metrics from intervention studies. (A)** Bacteriome and virome alpha diversity, **(B)** bacteriome beta diversity (weighted UniFrac distances), and **(C)** virome beta diversity (Bray-Curtis distances) from Experiment #2. **(D)** Bacteriome and virome alpha diversity, **(E)** bacteriome beta diversity (weighted UniFrac distances), and **(F)** virome beta diversity (Bray-Curtis distances) from Experiment #3. Alpha diversity trajectories were modelled using generalized additive mixed models (GAMMs). The fitted smooths represent the relationship between each metric and timing relative to first FVT dose (in days), with shaded ribbons indicating 95% confidence intervals. Greyed region corresponds to the time period where FVT doses were administered.

**Supplementary Figure 3. Relative abundance o**f differentially abundant taxa. Box plots representing the relative abundance of differentially abundant (A) bacterial taxa and (B) vOTUs from Experiment #2, and (C) vOTUs from Experiment #3. No bacterial taxa were significantly differentially abundant in Experiment #3.

**Supplementary Figure 4. Predicted replication cycles of vOTUs in the FVT.** Stacked bar charts displaying the proportion of FVT vOTUs assigned as temperate or non-temperate lifestyles in Experiments #2 and #3. vOTU counts (n) are indicated within the respective bars.

## References

1. Mertens A, Benjamin-Chung J, Colford JM, Hubbard AE, Van Der Laan MJ, Coyle J, Sofrygin O, Cai W, Jilek W, Rosete S, Nguyen A, Pokpongkiat NN, Djajadi S, Seth A, Jung E, Chung EO, Malenica I, Hejazi N, Li H, Hafen R, Subramoney V, Häggström J, Norman T, Christian P, Brown KH, Arnold BF, The Ki Child Growth Consortium, Ahmed T, Ali A, Begín F, Bessong PO, Bhutta ZA, Black RE, Bodhidatta L, Checkley W, Crabtree JE, Das R, Das S, Duggan CP, Faruque ASG, Fawzi WW, Da Silva Filho JQ, Gilman RH, Guerrant RL, Haque R, Houpt ER, Iqbal NT, John J, John SM, Kang G, Kosek M, Lima AÂM, Mahopo TC, Manandhar DS, Manji KP, Mduma E, Mohan VR, Moore SE, Nyathi ME, Olortegui MP, Petri WA, Premkumar PS, Prentice AM, Rahman N, Sadiq K, Sarkar R, Saville NM, Shrestha BP, Shrestha SK, Sonko B, Svensen E, Syed S, Umrani F, Ward HD, Yori PP. 2023. Child wasting and concurrent stunting in low- and middle-income countries. Nature 621:558–567.

2. Black RE, Victora CG, Walker SP, Bhutta ZA, Christian P, De Onis M, Ezzati M, Grantham-McGregor S, Katz J, Martorell R, Uauy R. 2013. Maternal and child undernutrition and overweight in low-income and middle-income countries. The Lancet 382:427–451.

3. United Nations Children’s Fund (UNICEF), World Health Organization, International Bank for Reconstruction and Development/The World Bank. 2025. Levels and trends in child malnutrition: UNICEF / WHO / World Bank Group Joint Child Malnutrition Estimates. Key findings of the 2025 edition. World Health Organization;, Geneva.

4. Subramanian S, Huq S, Yatsunenko T, Haque R, Mahfuz M, Alam MA, Benezra A, Destefano J, Meier MF, Muegge BD, Barratt MJ, VanArendonk LG, Zhang Q, Province MA, Petri WA, Ahmed T, Gordon JI. 2014. Persistent gut microbiota immaturity in malnourished Bangladeshi children. Nature 510:417–421.

5. Reyes A, Blanton LV, Cao S, Zhao G, Manary M, Trehan I, Smith MI, Wang D, Virgin HW, Rohwer F, Gordon JI. 2015. Gut DNA viromes of Malawian twins discordant for severe acute malnutrition. Proceedings of the National Academy of Sciences of the United States of America 112:11941–11946.

6. Robertson RC, Edens TJ, Carr L, Mutasa K, Gough EK, Evans C, Geum HM, Baharmand I, Gill SK, Ntozini R, Smith LE, Chasekwa B, Majo FD, Tavengwa NV, Mutasa B, Francis F, Tome J, Stoltzfus RJ, Humphrey JH, Prendergast AJ, Manges AR. 2023. The gut microbiome and early-life growth in a population with high prevalence of stunting. Nat Commun 14:654.

7. Vatanen T, Franzosa EA, Schwager R, Tripathi S, Arthur TD, Vehik K, Lernmark Å, Hagopian WA, Rewers MJ, She JX, Toppari J, Ziegler AG, Akolkar B, Krischer JP, Stewart CJ, Ajami NJ, Petrosino JF, Gevers D, Lähdesmäki H, Vlamakis H, Huttenhower C, Xavier RJ. 2018. The human gut microbiome in early-onset type 1 diabetes from the TEDDY study. Nature 562:589–594.

8. Zhao G, Vatanen T, Droit L, Park A, Kostic AD, Poon TW, Vlamakis H, Siljander H, Härkönen T, Hämäläinen AM, Peet A, Tillmann V, Ilonen J, Wang D, Knip M, Xavier RJ, Virgin HW. 2017. Intestinal virome changes precede autoimmunity in type I diabetes-susceptible children. Proceedings of the National Academy of Sciences of the United States of America 114:E6166–E6175.

9. Hoskinson C, Dai DLY, Del Bel KL, Becker AB, Moraes TJ, Mandhane PJ, Finlay BB, Simons E, Kozyrskyj AL, Azad MB, Subbarao P, Petersen C, Turvey SE. 2023. Delayed gut microbiota maturation in the first year of life is a hallmark of pediatric allergic disease. Nat Commun 14:4785.

10. Russell SL, Gold MJ, Hartmann M, Willing BP, Thorson L, Wlodarska M, Gill N, Blanchet MR, Mohn WW, McNagny KM, Finlay BB. 2012. Early life antibiotic-driven changes in microbiota enhance susceptibility to allergic asthma. EMBO Reports 13:440–447.

11. Smith MI, Yatsunenko T, Manary MJ, Trehan I, Mkakosya R, Cheng J, Kau AL, Rich SS, Concannon P, Mychaleckyj JC, Liu J, Houpt E, Li JV, Holmes E, Nicholson J, Knights D, Ursell LK, Knight R, Gordon JI. 2013. Gut Microbiomes of Malawian Twin Pairs Discordant for Kwashiorkor. Science 339:548–554.

12. Khan Mirzaei M, Khan MAA, Ghosh P, Taranu ZE, Taguer M, Ru J, Chowdhury R, Kabir MM, Deng L, Mondal D, Maurice CF. 2020. Bacteriophages isolated from stunted children can regulate gut bacterial communities in an age-specific manner. Cell Host and Microbe 27:199–212.

13. Khan Mirzaei M, Maurice CF. 2017. Ménage à trois in the human gut: interactions between host, bacteria and phages. Nature Reviews Microbiology 15:397–408.

14. Hsu BB, Gibson TE, Yeliseyev V, Liu Q, Lyon L, Bry L, Silver PA, Gerber GK. 2019. Dynamic Modulation of the Gut Microbiota and Metabolome by Bacteriophages in a Mouse Model. Cell Host and Microbe 25:803–814.e5.

15. Fortier L-C, Sekulovic O. 2013. Importance of prophages to evolution and virulence of bacterial pathogens. Virulence 4:354–365.

16. Rodriguez-Valera F, Martin-Cuadrado A-B, Rodriguez-Brito B, Pašić L, Thingstad TF, Rohwer F, Mira A. 2009. Explaining microbial population genomics through phage predation. Nat Rev Microbiol 7:828–836.

17. Chevallereau A, Pons BJ, Van Houte S, Westra ER. 2022. Interactions between bacterial and phage communities in natural environments. Nat Rev Microbiol 20:49–62.

18. Shamash M, Maurice CF. 2026. A meta-analysis of infant gut viromes reveals global patterns in bacteriophage community assembly and functional capacity over the first three years of life. Nat Commun 17:7698.

19. Sinha A, Li Y, Mirzaei MK, Shamash M, Samadfam R, King IL, Maurice CF. 2022. Transplantation of bacteriophages from ulcerative colitis patients shifts the gut bacteriome and exacerbates the severity of DSS colitis. Microbiome 10:105.

20. Brunse A, Deng L, Pan X, Hui Y, Castro-Mejía JL, Kot W, Nguyen DN, Secher JB-M, Nielsen DS, Thymann T. 2022. Fecal filtrate transplantation protects against necrotizing enterocolitis. ISME J 16:686–694.

21. Spiegelhauer MR, Offersen SM, Mao X, Gambino M, Sandris Nielsen D, Nguyen DN, Brunse A. 2025. Protection against experimental necrotizing enterocolitis by fecal filtrate transfer requires an active donor virome. Gut Microbes 17:2486517.

22. Offersen SM, Mao X, Spiegelhauer MR, Larsen F, Li VR, Sandris Nielsen D, Aunsholt L, Thymann T, Brunse A. 2024. Fecal virus-like particles are sufficient to reduce necrotizing enterocolitis. Gut Microbes 16:2392876.

23. Shamash M, Maurice CF. 2021. Phages in the infant gut: a framework for virome development during early life. The ISME Journal 16:323–330.

24. Fahur Bottino G, Bonham KS, Patel F, McCann S, Zieff M, Naspolini N, Ho D, Portlock T, Joos R, Midani FS, Schüroff P, Das A, Shennon I, Wilson BC, O’Sullivan JM, Britton RA, Murray DM, Kiely ME, Taddei CR, Beltrão-Braga PCB, Campos AC, Polanczyk GV, Huttenhower C, Donald KA, Klepac-Ceraj V. 2025. Early life microbial succession in the gut follows common patterns in humans across the globe. Nat Commun 16:660.

25. Brown EM, Wlodarska M, Willing BP, Vonaesch P, Han J, Reynolds LA, Arrieta M-CC, Uhrig M, Scholz R, Partida O, Borchers CH, Sansonetti PJ, Finlay BB. 2015. Diet and specific microbial exposure trigger features of environmental enteropathy in a novel murine model. Nature communications 6:7806.

26. Huus KE, Bauer KC, Brown EM, Bozorgmehr T, Woodward SE, Serapio-Palacios A, Boutin RCT, Petersen C, Finlay BB. 2020. Commensal Bacteria Modulate Immunoglobulin A Binding in Response to Host Nutrition. Cell Host and Microbe 27:909–921.e5.

27. Bhattacharjee A, Burr AHP, Overacre-Delgoffe AE, Tometich JT, Yang D, Huckestein BR, Linehan JL, Spencer SP, Hall JA, Harrison OJ, Morais da Fonseca D, Norton EB, Belkaid Y, Hand TW. 2021. Environmental enteric dysfunction induces regulatory T cells that inhibit local CD4+ T cell responses and impair oral vaccine efficacy. Immunity 54:1745–1757.e7.

28. Bauer KC, York EM, Cirstea MS, Radisavljevic N, Petersen C, Huus KE, Brown EM, Bozorgmehr T, Berdún R, Bernier L, Lee AHY, Woodward SE, Krekhno Z, Han J, Hancock REW, Ayala V, MacVicar BA, Finlay BB. 2022. Gut microbes shape microglia and cognitive function during malnutrition. Glia 70:820–841.

29. £ypaczewski P, Shapiro BJ. Dibenzocyclooctyne-modified PCR primers enable direct enzyme-free click chemistry ligation for custom nanopore amplicon sequencing.

30. Curry KD, Wang Q, Nute MG, Tyshaieva A, Reeves E, Soriano S, Wu Q, Graeber E, Finzer P, Mendling W, Savidge T, Villapol S, Dilthey A, Treangen TJ. 2022. Emu: species-level microbial community profiling of full-length 16S rRNA Oxford Nanopore sequencing data. Nat Methods 19:845–853.

31. McMurdie PJ, Holmes S. 2013. phyloseq: An R Package for Reproducible Interactive Analysis and Graphics of Microbiome Census Data. PLoS ONE 8:e61217.

32. Oksanen J, Blanchet FG, Friendly M, Kindt R, Legendre P, McGlinn D, Minchin PR, O’Hara RB, Simpson GL, Solymos P, Stevens MHH, Szoecs E, Wagner H. 2019. vegan: Community Ecology Package.

33. Liland KH, Mevik B-H, Wehrens R. 2024. pls: Partial Least Squares and Principal Component Regression. https://CRAN.R-project.org/package=pls.

34. Wood SN. 2011. Fast Stable Restricted Maximum Likelihood and Marginal Likelihood Estimation of Semiparametric Generalized Linear Models. Journal of the Royal Statistical Society Series B: Statistical Methodology 73:3–36.

35. Lenth RV, Piaskowski J. 2025. emmeans: Estimated Marginal Means, aka Least-Squares Means. https://rvlenth.github.io/emmeans/.

36. Gu Z, Eils R, Schlesner M. 2016. Complex heatmaps reveal patterns and correlations in multidimensional genomic data. Bioinformatics 32:2847–2849.

37. Shamash M, Kapoor S, Maurice CF. 2025. Benchmarking of a time-saving and scalable protocol for the extraction of DNA from diverse viromes. PeerJ 13:e18785.

38. Chen S, Zhou Y, Chen Y, Gu J. 2018. fastp: an ultra-fast all-in-one FASTQ preprocessor. Bioinformatics 34:i884–i890.

39. Langmead B, Salzberg SL. 2012. Fast gapped-read alignment with Bowtie 2. Nature Methods 9:357–359.

40. Bankevich A, Nurk S, Antipov D, Gurevich AA, Dvorkin M, Kulikov AS, Lesin VM, Nikolenko SI, Pham S, Prjibelski AD, Pyshkin AV, Sirotkin AV, Vyahhi N, Tesler G, Alekseyev MA, Pevzner PA. 2012. SPAdes: a new genome assembly algorithm and its applications to single-cell sequencing. Journal of computational biology : a journal of computational molecular cell biology, 2012/04/16 ed. 19:455–477.

41. Nurk S, Meleshko D, Korobeynikov A, Pevzner PA. 2017. metaSPAdes: a new versatile metagenomic assembler. Genome Res 27:824–834.

42. Camargo AP, Roux S, Schulz F, Babinski M, Xu Y, Hu B, Chain PSG, Nayfach S, Kyrpides NC. 2024. Identification of mobile genetic elements with geNomad. Nat Biotechnol 42:1303–1312.

43. Camacho C, Coulouris G, Avagyan V, Ma N, Papadopoulos J, Bealer K, Madden TL. 2009. BLAST+: architecture and applications. BMC Bioinformatics 10:421.

44. Nayfach S, Camargo AP, Schulz F, Eloe-fadrosh E, Roux S, Kyrpides NC. CheckV assesses the quality and completeness of metagenome-assembled viral genomes. Nature Biotechnology 39:578–585.

45. Li H, Handsaker B, Wysoker A, Fennell T, Ruan J, Homer N, Marth G, Abecasis G, Durbin R. 2009. The Sequence Alignment/Map format and SAMtools. Bioinformatics 25:2078–2079.

46. Roux S, Emerson JB, Eloe-Fadrosh EA, Sullivan MB. 2017. Benchmarking viromics: An in silico evaluation of metagenome-enabled estimates of viral community composition and diversity. PeerJ 2017:e3817.

47. Roux S, Camargo AP, Coutinho FH, Dabdoub SM, Dutilh BE, Nayfach S, Tritt A. 2023. iPHoP: An integrated machine learning framework to maximize host prediction for metagenome-derived viruses of archaea and bacteria. PLoS Biol 21:e3002083.

48. Hockenberry AJ, Wilke CO. 2021. BACPHLIP: predicting bacteriophage lifestyle from conserved protein domains. PeerJ 9:e11396.

49. R Core Team. 2020. R: A Language and Environment for Statistical Computing. R Foundation for Statistical Computing, Vienna, Austria.

50. Simpson GL. 2024. gratia: An R package for exploring generalized additive models. Journal of Open Source Software 9:6962.

51. Nickols WA, Kuntz T, Shen J, Maharjan S, Mallick H, Franzosa EA, Thompson KN, Nearing JT, Huttenhower C. 2026. MaAsLin 3: Refining and extending generalized multivariable linear models for meta-omic association discovery. Nature Methods 23:554–564.

52. Walter J, Armet AM, Finlay BB, Shanahan F. 2020. Establishing or Exaggerating Causality for the Gut Microbiome: Lessons from Human Microbiota-Associated Rodents. Cell 180:221–232.

53. Fouladi F, Glenny EM, Bulik-Sullivan EC, Tsilimigras MCB, Sioda M, Thomas SA, Wang Y, Djukic Z, Tang Q, Tarantino LM, Bulik CM, Fodor AA, Carroll IM. 2020. Sequence variant analysis reveals poor correlations in microbial taxonomic abundance between humans and mice after gnotobiotic transfer. The ISME Journal 14:1809–1820.

54. Li Y, Cao W, Gao NL, Zhao X-M, Chen W-H. 2022. Consistent Alterations of Human Fecal Microbes After Transplantation into Germ-Free Mice. Genomics, Proteomics & Bioinformatics 20:382–393.

55. Stewart CJ, Ajami NJ, O’Brien JL, Hutchinson DS, Smith DP, Wong MC, Ross MC, Lloyd RE, Doddapaneni HV, Metcalf GA, Muzny D, Gibbs RA, Vatanen T, Huttenhower C, Xavier RJ, Rewers M, Hagopian W, Toppari J, Ziegler AG, She JX, Akolkar B, Lernmark A, Hyoty H, Vehik K, Krischer JP, Petrosino JF. 2018. Temporal development of the gut microbiome in early childhood from the TEDDY study. Nature 562:583–588.

56. De Sordi L, Lourenço M, Debarbieux L. 2019. The Battle Within: Interactions of Bacteriophages and Bacteria in the Gastrointestinal Tract. Cell Host & Microbe 25:210–218.

57. Yan Y, Nguyen LH, Franzosa EA, Huttenhower C. 2020. Strain-level epidemiology of microbial communities and the human microbiome. Genome Med 12:71.

58. Middelboe M, Holmfeldt K, Riemann L, Nybroe O, Haaber J. 2009. Bacteriophages drive strain diversification in a marine *Flavobacterium* : implications for phage resistance and physiological properties. Environmental Microbiology 11:1971–1982.

59. Rodriguez-Valera F, Martín-Cuadrado A-B, Rodriguez-Brito B, Pasic L, Thingstad TF, Rohwer F, Mira A. 2009. Explaining microbial population genomics through phage predation. Nat Prec 10.1038/npre.2009.3489.1.

60. Bondy-Denomy J, Davidson AR. 2014. When a virus is not a parasite: the beneficial effects of prophages on bacterial fitness. J Microbiol 52:235–242.

61. Chen RY, Mostafa I, Hibberd MC, Das S, Mahfuz M, Naila NN, Islam MM, Huq S, Alam MA, Zaman MU, Raman AS, Webber D, Zhou C, Sundaresan V, Ahsan K, Meier MF, Barratt MJ, Ahmed T, Gordon JI. 2021. A Microbiota-Directed Food Intervention for Undernourished Children. New England Journal of Medicine 384.

62. Chen RY, Mostafa I, Hibberd MC, Das S, Lynn HM, Webber DM, Mahfuz M, Barratt MJ, Ahmed T, Gordon JI. 2021. Melding microbiome and nutritional science with early child development. Nat Med 27:1503–1506.

