## Supplementary figures and images for "A human microbiota-associated mouse model of early-life malnutrition reveals persistent microbiome immaturity and limited response to fecal viral transplantation"

### Supplementary Figure 1

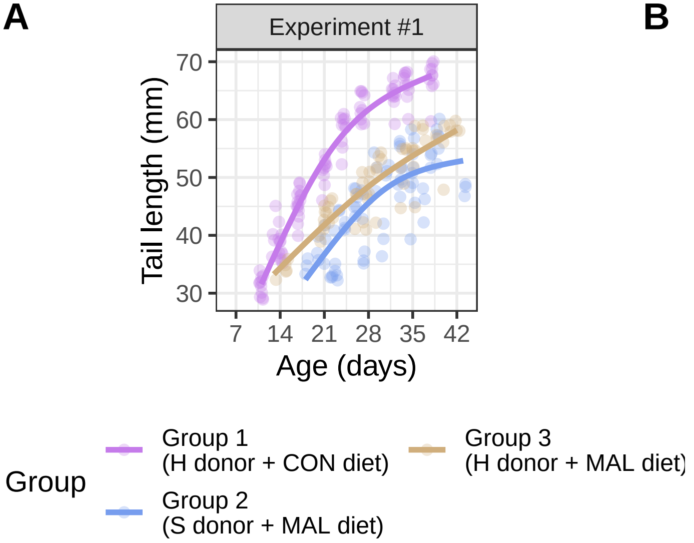

### Supplementary Figure 2

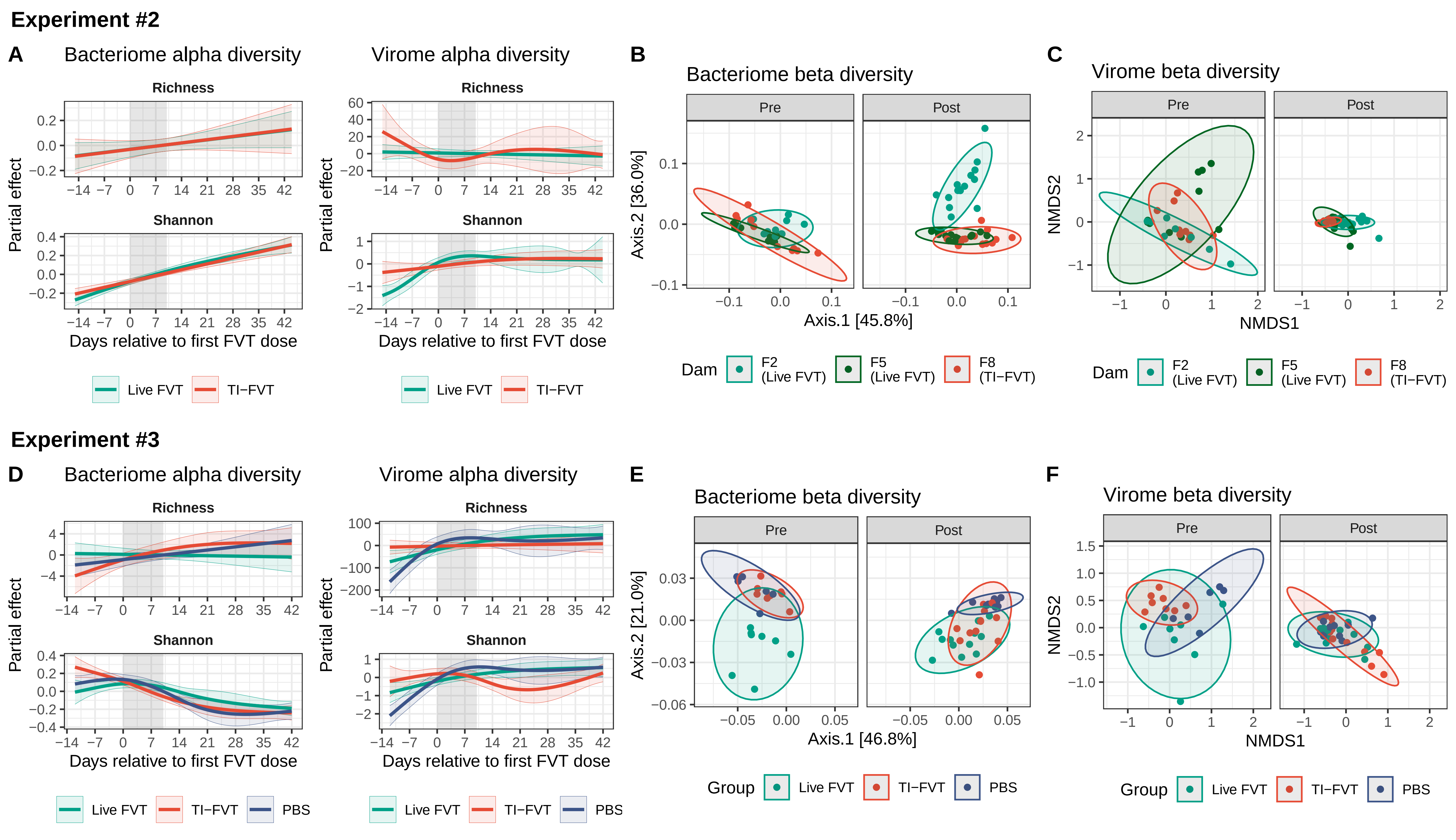

### Supplementary Figure 3

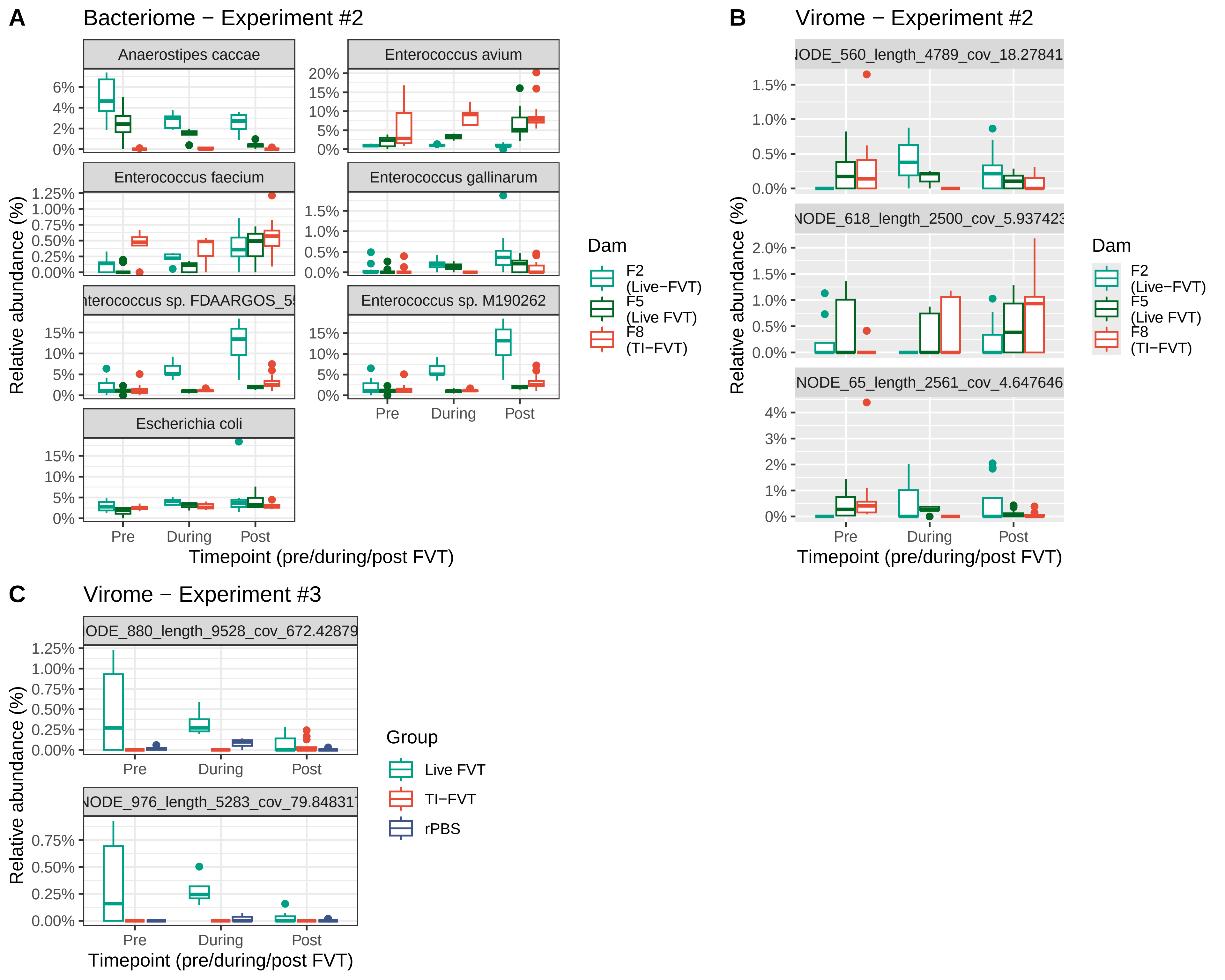

### Supplementary Figure 4

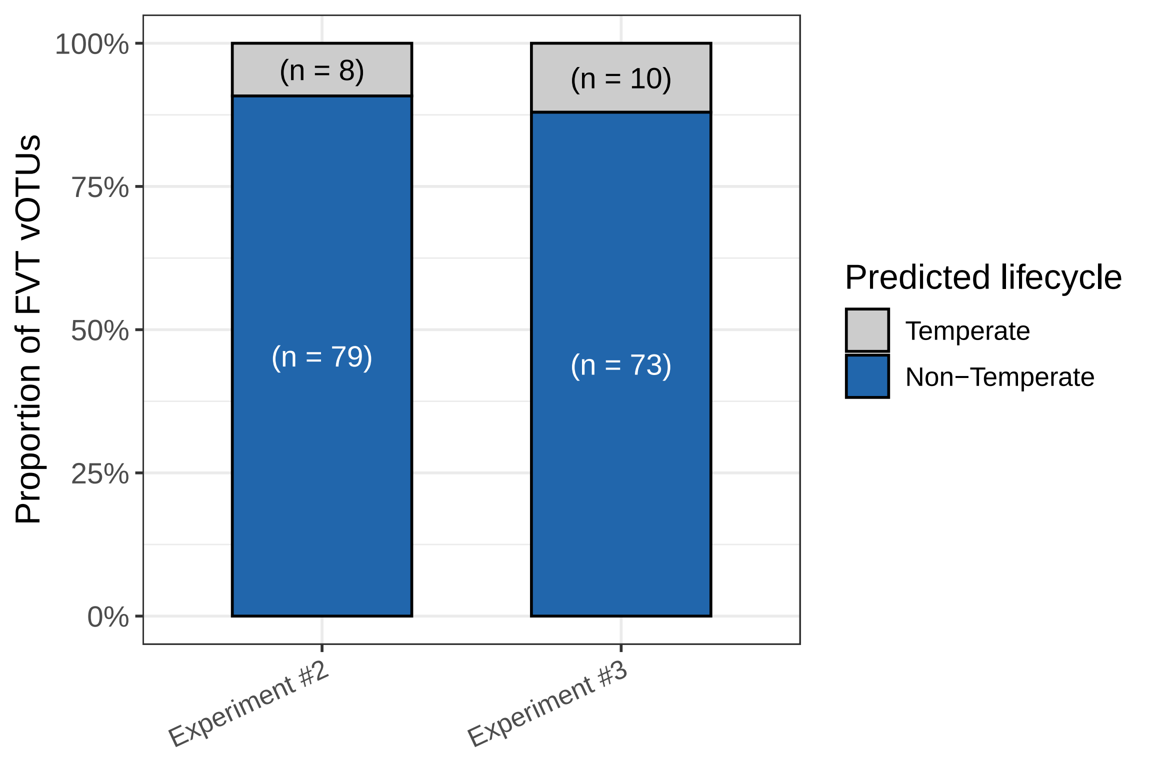
